# Investigating the reliability of molecular estimates of evolutionary time when substitution rates and speciation rates vary

**DOI:** 10.1101/2021.06.27.450013

**Authors:** Andrew M. Ritchie, Xia Hua, Lindell Bromham

## Abstract

**Background:** An accurate timescale of evolutionary history is essential to testing hypotheses about the influence of historical events and processes, and the timescale for evolution is increasingly derived from analysis of DNA sequences. But variation in the rate of molecular evolution complicates the inference of time from DNA. Evidence is growing for numerous factors, such as life history and habitat, that are linked both to the molecular processes of mutation and fixation and to rates of macroevolutionary diversification. However, the most widely used methods rely on idealised models of rate variation, such as the uncorrelated and autocorrelated clocks, and molecular dating methods are rarely tested against complex models of rate change. One relationship that is not accounted for in molecular dating is the potential for interaction between molecular substitution rates and speciation, a relationship that has been supported by empirical studies in a growing number of taxa. If these relationships are as widespread as current evidence suggests, they may have a significant influence on molecular dates.

**Results:** We simulate phylogenies and molecular sequences under three different realistic rate variation models – one in which speciation rates and substitution rates both vary but are unlinked, one in which they covary continuously and one punctuated model in which molecular change is concentrated in speciation events, using empirical case studies to parameterise realistic simulations. We test three commonly used “relaxed clock” molecular dating methods against these realistic simulations to explore the degree of error in molecular dates under each model. We find average divergence time inference errors ranging from 12% of node age for the unlinked model when reconstructed under an uncorrelated rate prior using BEAST 2, to up to 91% when sequences evolved under the punctuated model are reconstructed under an autocorrelated prior using PAML.

**Conclusions:** We demonstrate the potential for substantial errors in molecular dates when both speciation rates and substitution rates vary between lineages. This study highlights the need for tests of molecular dating methods against realistic models of rate variation generated from empirical parameters and known relationships.

## 1. Background

Understanding the timescale of evolution is critical to researching the processes that generate and shape the diversity of life on Earth. Analysis of DNA sequences offers the possibility of investigating the tempo and mode of evolution of all extant lineages, not just those that have a detailed fossil record. The earliest molecular dating analyses concluded that the average rate of molecular evolution was relatively constant over time in some proteins because of an apparently linear accumulation of amino acid differences when compared between species (1, 2). However, it has since become apparent that the average rate of molecular evolution can be influenced by a wide range of species traits, such as generation time, body size and longevity (3, 4), by macroevolutionary processes such as net diversification rate (5–10) and potentially also by environmental features like temperature (9–11). If the average rate of molecular evolution varies substantially and consistently between lineages, then we may need to incorporate this rate variation into models of molecular evolution in order to accurately infer divergence times among species (12).

A wide variety of models have been developed to account for different patterns of among-lineage molecular rate variation (13–16). In the first instance, these models differ in how many different evolutionary rate parameters are estimated for different parts of the phylogeny. For example, a ‘strict clock’ model estimates only one rate parameter that applies to every branch in the tree, while a ‘local clock’ model estimates additional rate parameters for subsets of branches (17–19), and a ‘relaxed clock’ provides for a separate rate on every branch (20). Models also differ in how the rates on different branches are related to one another. Some models, notably in the penalized-likelihood framework, aim to minimize the overall variation of branch-specific rates or the differences between adjacent branches (21, 22). For Bayesian phylogenetic methods (23, 24), branch-specific rates are drawn from a prior probability distribution that determines which rates and patterns of relative rate variation are considered *a priori* more probable (we refer to this prior as the rate prior). Common variants include uncorrelated models, in which each branch-specific rate is an independent draw from a distribution with a common mean (20), and autocorrelated models, in which the mean of the distribution for each branch-specific rate depends on the rate drawn for its parent branch (25, 26).

Testing over the last two decades has generally supported the robustness of divergence time estimates under the Bayesian methodology to the choice of rate prior, provided that enough separate rate parameters are allowed and sufficient calibrating information is provided (27–33); but see (34). However, these studies rely heavily on simulated data or on specific and usually well-calibrated empirical examples, whereas simulations have indicated that different rate priors may give different results when calibrating information is poorer (30). Changing the rate prior in these circumstances could lead to differences in empirical dates and associated causal hypotheses. Examples of empirical studies where the choice of rate prior appears to be a factor in differing age estimates include influenza viruses (35), the age of grasstrees (36) and the origins of Metazoa (37).

Currently used models of rate variation are generally either uncorrelated, so that each lineage-specific rate is drawn independently from a common distribution, or autocorrelated so that lineage-specific rates evolve from the rates of their parent lineages according to a stochastic, random walk. However, the factors shaping variation in rate of molecular evolution are complex: in addition to random variation in rates between lineages, there is clearly also systematic variation influenced by many factors including species traits, population dynamics, environment and evolutionary history (38). Evolutionary rates can be faster at lower elevations and latitudes (9, 39, 40) but slower in arid climates (41). Life history can covary with molecular rates across multiple axes, including body size (42–45), fecundity (46), generation time (47) and longevity (48, 49). Associations have also been observed between substitution rates and more complex species traits, such as parasitism (50), flightlessness (51) and sexual competition (52). Furthermore, substitution rates are the result of underlying microevolutionary processes, and may therefore be affected by population size (53–56).

In addition to covarying with other evolving traits, there is evidence that variation in molecular rates can be associated with macroevolutionary patterns. Relationships between the relative size of sister clades, representing their net diversification rates, and their molecular rates have been detected in several studies, include in reptiles (8, 57), birds (7, 58), flowering plants (6, 59, 60) and across Metazoa (61). Several studies have also noted relationships between the genetic divergence of species in a phylogeny and the number of intervening speciation events, which may also represent examples of this phenomenon (62–64).

The mechanism behind observed correlations between diversification and molecular evolution remains a matter of debate. One line of argument proposes that higher molecular rates could promote speciation by speeding the development of reproductive isolation (65). In this hypothesis, subpopulations become sporadically isolated over time and may accumulate alleles that may be effectively neutral on their own but deleterious in combination (66, 67). In this case, greater rates of molecular evolution could lead to faster development of reproductive isolation during periods of isolation, leading to a greater average rate of speciation (68, 69). Alternatively, higher substitution rates could reduce extinction rates by accumulating more standing genetic diversity which provides a buffer against changing environment by allowing more rapid selective response (70). Speciation rates and molecular rates could also be related through a common driving factor, such as climate or environmental productivity (5, 10). In the other causal direction, speciation itself could increase average rates of molecular evolution by producing rapid bursts of evolution (62, 63, 71). Suggested mechanisms for an increase in substitution rates during speciation include an increase in fixation rates of mildly deleterious alleles and linked neutral mutations due to drift, or large-scale genomic divergence caused by an increase in the frequency of disruptive genomic events such as polyploidy or chromosomal rearrangements (71).

An association between substitution rates and diversification rates could affect the performance of existing divergence time estimation methods (e.g. 72). In particular, Bayesian methods divide their specification of prior beliefs about the evolutionary process between the rate prior, the prior on divergence times and tree topology (sometimes referred to as the tree prior or node time prior), and calibrating distributions on the ages of individual speciation events (23, 73). These priors are essential to estimating divergence time, because genetic distances between sequences are the product of rate and time (the total number of character differences between related sequences). This means that even if genetic distances are correctly inferred on a phylogeny, rate and time are unidentifiable without additional constraints on joint values they can hold (74, 75). The rate prior, node time prior and calibrating distributions place either soft or hard constraints on these values, allowing researchers to infer evolutionary history on an absolute timescale. However, this means that inferred divergence times are always subject to the choice of rate, time and calibration priors even with large amounts of sequence data (76, 77). Most commonly used Bayesian molecular dating methods do not propose *a priori* relationships between the rate prior and the tree prior, so the joint distribution implied by these priors will assign relatively higher probability to l long branches with high substitution rates and short branches with low substitution rates than would be expected when rates and times are related. It is therefore possible that an unmodelled relationship between rates and times could produce distributions of molecular rates and speciation times that differ substantially from those implied by common Bayesian priors, leading to widespread error in estimates of evolutionary dates.

Here, we investigate the potential influence of speciation-related rates of molecular evolution on the reliability of molecular dating studies. Rather than assuming only one specific model of the association between diversification rates and rates of molecular evolution, we model a range of possible causal links. We include three alternative models of the association between rates of molecular evolution and speciation rates. In the first, both speciation rates and substitution rates vary over the phylogeny, but neither influences the other. In the second, we model continuous variation in a linked manner, which might represent either direct influence of one on the other (65) or an indirect association between speciation rates and substitution rates, for example if they are both influenced by environmental factors (5, 10). The third model is a punctuated model in which bursts of substitution are associated with speciation events (62, 63, 71, 78). The proportion of all substitutions associated with such bursts has recently been shown to impact Bayesian inference of clade crown ages under an uncorrelated lognormal rate prior (72).

In this study, we simulate the evolution of molecular sequences under all three models of the link between speciation and rate of molecular evolution – unlinked, continuous coevolution and punctuated models – and observe the average effects on errors in Bayesian divergence time estimates. Most simulation studies use arbitrarily chosen values to parameterize the simulations. In order to make our test have real-world significance, we base our simulations, as far as possible, on empirically determined relationships between rates of molecular evolution and diversification rate. As a convenient well-studied case study, we parameterize our models using values taken from empirical inferences on bird data (79, 80), or by using parameter values which reproduce established empirical relationships between rates of molecular evolution and diversification rate in birds (81). We have chosen birds as a convenient exemplar since an association between rates of molecular evolution and rates of diversification has been reported for birds (58), and there are few if any other taxa whose rates of molecular evolution and diversification have been as well-described. We test the performance of three commonly used “relaxed clock” molecular dating methods: since the choice of rate prior and methodological details may affect the results of phylogeny reconstruction (34), we have tested uncorrelated and autocorrelated rate models in two different molecular dating programs. We reconstruct divergence times on these simulated data sets and examine the effects of different forms of correlated molecular and speciation rates on the performance of molecular divergence time reconstruction.

## 2. Results

### 2.1 Validation of tree simulations

In order to ensure our simulations are as close as possible to real-world data, we chose to parameterize the simulations using empirical estimates of parameters from bird data, because birds are one of the most well-studied groups for both substitution rate variation and diversification rate. While these parameters might not exactly capture processes in other taxa, we believe that basing the simulations on a real case study lends a degree of realism that would be lacking if we arbitrarily chose convenient parameter values. We checked that the simulation procedure produced trees with average characteristics similar to the avian clades used to parameterize the study (80 [supplementary data], 82). Across all data sets, the mean root date of simulated trees was 28.4 million years, compared to a range of 23.8-35.6 million years for bird clades with 70-80 taxa. Simulated lineage-specific speciation rates at the tips were on average slightly lower than the speciation rate of similar clades in the bird tree (80). The range of simulated speciation rates on log scale was 0.009-0.92 compared to a range of 0.034-2.25 average log lineages/million years for bird clades with 70-80 taxa, similar taxon numbers to our simulated trees. The range of speciation rates within each tree was also similar to the range across the avian clades with similar taxon numbers, with an average magnitude of 0.44 log lineages/million years compared to 0.32-2.21 for avian clades with 70- 80 taxa. These observations suggest that the simulations produce realistic phylogenies, similar to those associated with real taxa.

### 2.2 Error in reconstructed trees

Trees were simulated under three models: Unlinked (speciation rates and molecular rates vary independently), Continuous (speciation rates and molecular rates covary), and Punctuated (molecular change occurs in bursts associated with speciation events as well as continuously within lineages) (Table 1). We used three different metrics, plus the gamma statistic and a sister pair analysis, to assess the accuracy of the reconstruction of trees simulated from the Unlinked model, the Continuous model, and the Punctuated model, under uncorrelated lognormal (UCLN) and autocorrelated lognormal (ACLN) models of among-lineage rate variation (Table 2).

**Table 1.** Simulation design.

| Simulation type | Speciation rates covary with substitution rates? | Brownian motion covariance | Punctuated burst of substitutions at nodes? |
| --- | --- | --- | --- |
| *Unlinked* | *No* | 0 | *no* |
| *Continuous* | *Yes* | 0.0044 | *no* |
| *Punctuated* | *No* | 0 | *yes* |

We lay out the conditions for each set of simulated trees. Each set consisted of 50 trees. Lineage-specific speciation rates and substitution rates evolve continuously through time via Brownian motion in all three studies. The three simulations differ in whether the covariance between speciation and substitution rates is greater than zero in the Brownian motion and whether an additional burst of substitutions occurs at nodes of the tree (speciation events).

**Table 2.**
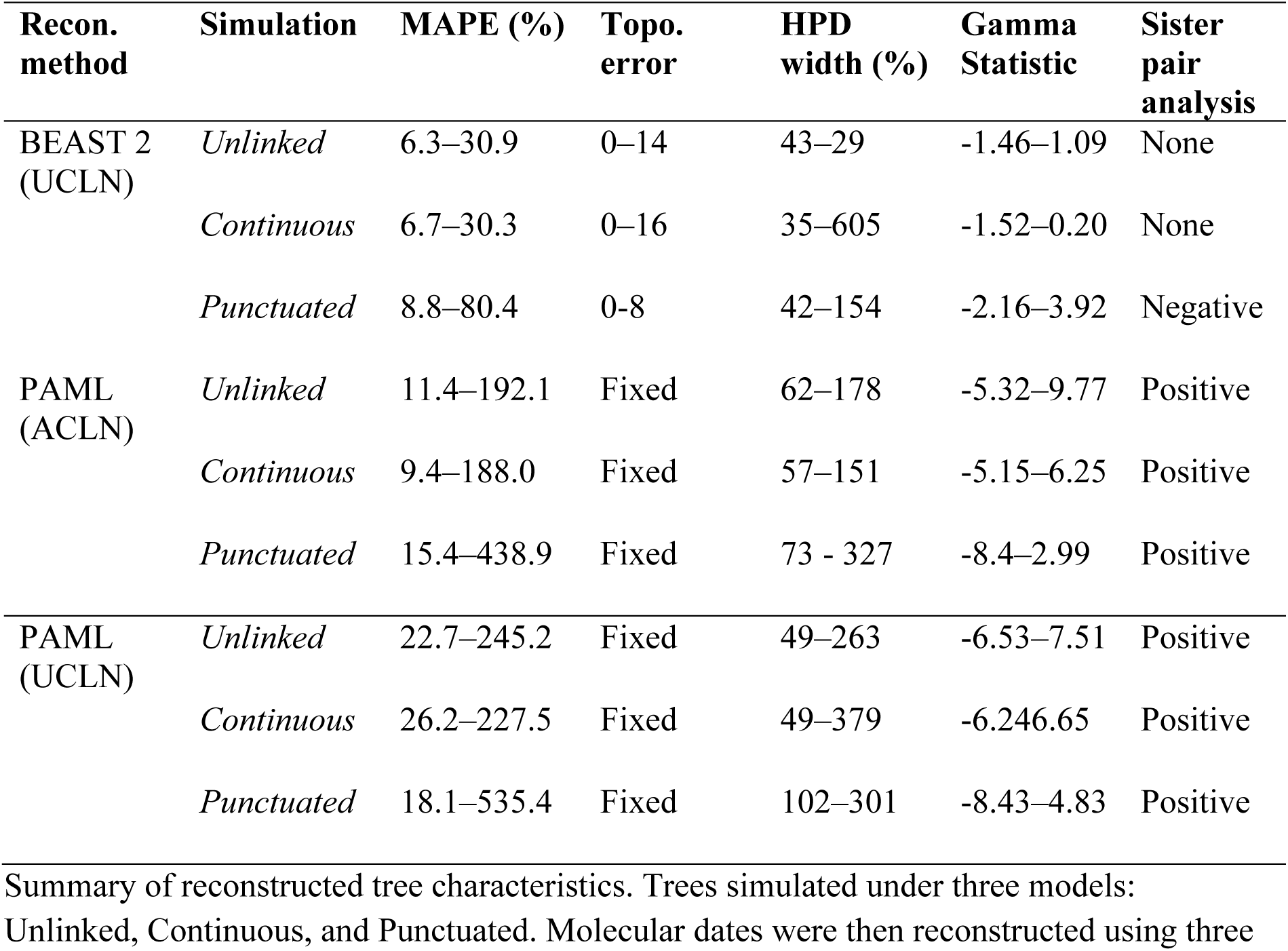
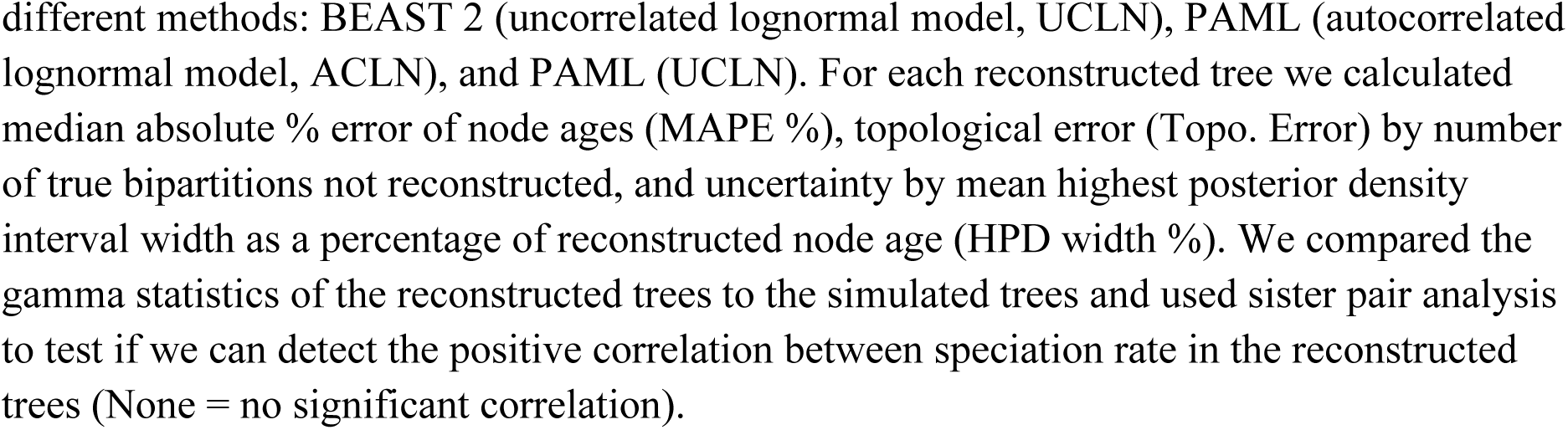
Summary of results.

| Recon. method | Simulation | MAPE (%) | Topo. error | HPD width (%) | Gamma Statistic | Sister pair analysis |
| --- | --- | --- | --- | --- | --- | --- |
| BEAST 2 (UCLN) | *Unlinked* | 6.3–30.9 | 0–14 | 43–29 | -1.46–1.09 | None |
|  | *Continuous* | 6.7–30.3 | 0–16 | 35–605 | -1.52–0.20 | None |
|  | *Punctuated* | 8.8–80.4 | 0–8 | 42–154 | -2.16–3.92 | Negative |
| PAML (ACLN) | *Unlinked* | 11.4–192.1 | Fixed | 62–178 | -5.32–9.77 | Positive |
|  | *Continuous* | 9.4–188.0 | Fixed | 57–151 | -5.15–6.25 | Positive |
|  | *Punctuated* | 15.4–438.9 | Fixed | 73 - 327 | -8.4–2.99 | Positive |
| PAML (UCLN) | *Unlinked* | 22.7–245.2 | Fixed | 49–263 | -6.53–7.51 | Positive |
|  | *Continuous* | 26.2–227.5 | Fixed | 49–379 | -6.246.65 | Positive |
|  | *Punctuated* | 18.1–535.4 | Fixed | 102–301 | -8.43–4.83 | Positive |

Summary of reconstructed tree characteristics. Trees simulated under three models: Unlinked, Continuous, and Punctuated. Molecular dates were then reconstructed using three
different methods: BEAST 2 (uncorrelated lognormal model, UCLN), PAML (autocorrelated lognormal model, ACLN), and PAML (UCLN). For each reconstructed tree we calculated median absolute % error of node ages (MAPE %), topological error (Topo. Error) by number of true bipartitions not reconstructed, and uncertainty by mean highest posterior density interval width as a percentage of reconstructed node age (HPD width %). We compared the gamma statistics of the reconstructed trees to the simulated trees and used sister pair analysis to test if we can detect the positive correlation between speciation rate in the reconstructed trees (None = no significant correlation).

Node dates were then inferred using three different reconstruction methods: BEAST 2 (83, 84) using the uncorrelated lognormal rate prior (UCLN); PAML (85) using the autocorrelated lognormal rate prior (ACLN); and PAML using UCLN. To compare the reconstructed molecular dates to the true dates from the simulated data, we calculated three metrics for each tree and summarised these over each simulation type (Table 2). The metrics were median absolute percentage error of node ages (MAPE), topological error (Topo. Error) by number of true bipartitions not reconstructed, and uncertainty by mean highest posterior density interval width as a percentage of reconstructed node age (HPD width %). We compared the gamma statistics of the reconstructed trees to the simulated trees in order to detect any systematic bias in node age error. As a way of evaluating the degree to which the modelled relationship between speciation and substitution rates is reflected in the phylogenies reconstructed from the simulated data, we also performed sister pair analysis to test if we can detect the positive correlation between speciation rate and substitution rate from the reconstructed trees.

First, we examined the median absolute percentage error (MAPE) in the ages of all nodes within each tree whose topology was correctly reconstructed (Figure 2). For trees reconstructed using BEAST 2 (UCLN), the average (min–max) MAPE was 12.2% for Unlinked simulations (6.3–30.9%). Errors for the Continuous simulation were similar at 14.2% (6.7–30.3%). Average MAPE for the Punctuated simulation was higher than for Unlinked trees at 20.3% (8.8–80.4%). For trees reconstructed using PAML (ACLN), MAPE values were higher and range of errors were larger than for BEAST 2 (UCLN). Average MAPE for Unlinked trees was 66.0% (11–192%), compared to 60.6% (9.4%–188%) for Continuous trees and 91.1% (15.4–438.9%) for Punctuated trees. For reconstructions with PAML (UCLN), average MAPE was even higher than PAML (ACLN) at 77.0% (22.7– 245.2) for Unlinked trees, for Continuous trees 69.6% (26.2–227.5), but lower for Punctuated trees at 74.9% (18.1-535.3). Because these errors can only be calculated for nodes that are shared between the true and reconstructed trees, we also examined an alternative error metric using the branch score (Kühner-Felsenstein) distance, which is based on the sum of squared branch length errors, includes all nodes, and treats branches that are incorrectly reconstructed as length zero, thus accounting for topology errors (86). This metric led to similar patterns and is included as online Supplementary Information (Figure S1).

**Figure 2.**
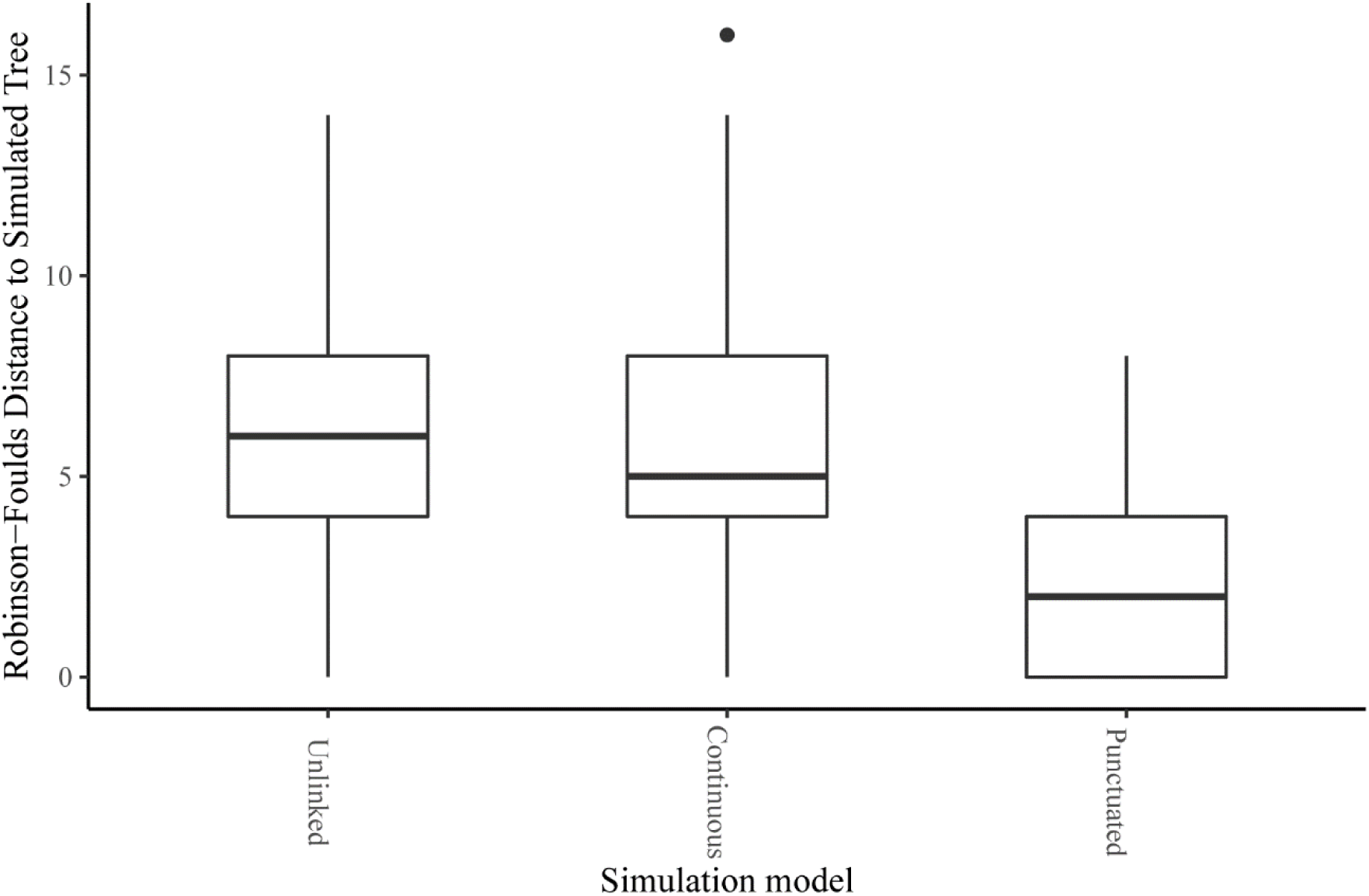
Topological error of trees reconstructed from data sets simulated under three simulation types with different relationships between molecular evolution and speciation (see Table 1). Topological error is given by the Robinson-Foulds distance (number of non-shared bipartitions). The three simulation models are Unlinked (instantaneous covariance of molecular rates and speciation rates = 0), Continuous (instantaneous covariance = 0.0044), and Punctuated (instantaneous covariance = 0, bursts of substitutions added at speciation events). Topologies and node times were reconstructed using the uncorrelated lognormal ‘relaxed clock’ rate prior in BEAST 2 (UCLN). No results are available for PAML because the topology was fixed for these analyses.

Second, for trees reconstructed using BEAST 2 (UCLN), we also investigated the degree of topological error via the Robinson-Foulds distance between the true and estimated tree (87). This measure can only be used for BEAST 2 because the ‘mcmctree’ program in PAML does not infer topology. The median background topological error for the Unlinked simulations was 6 out of 74 bipartitions (0–14), compared to 5 (0–16) for the Continuous model (Figure 3). Error was lower for the Punctuated model with a median distance of 2 bipartitions (0–8).

**Figure 3.**
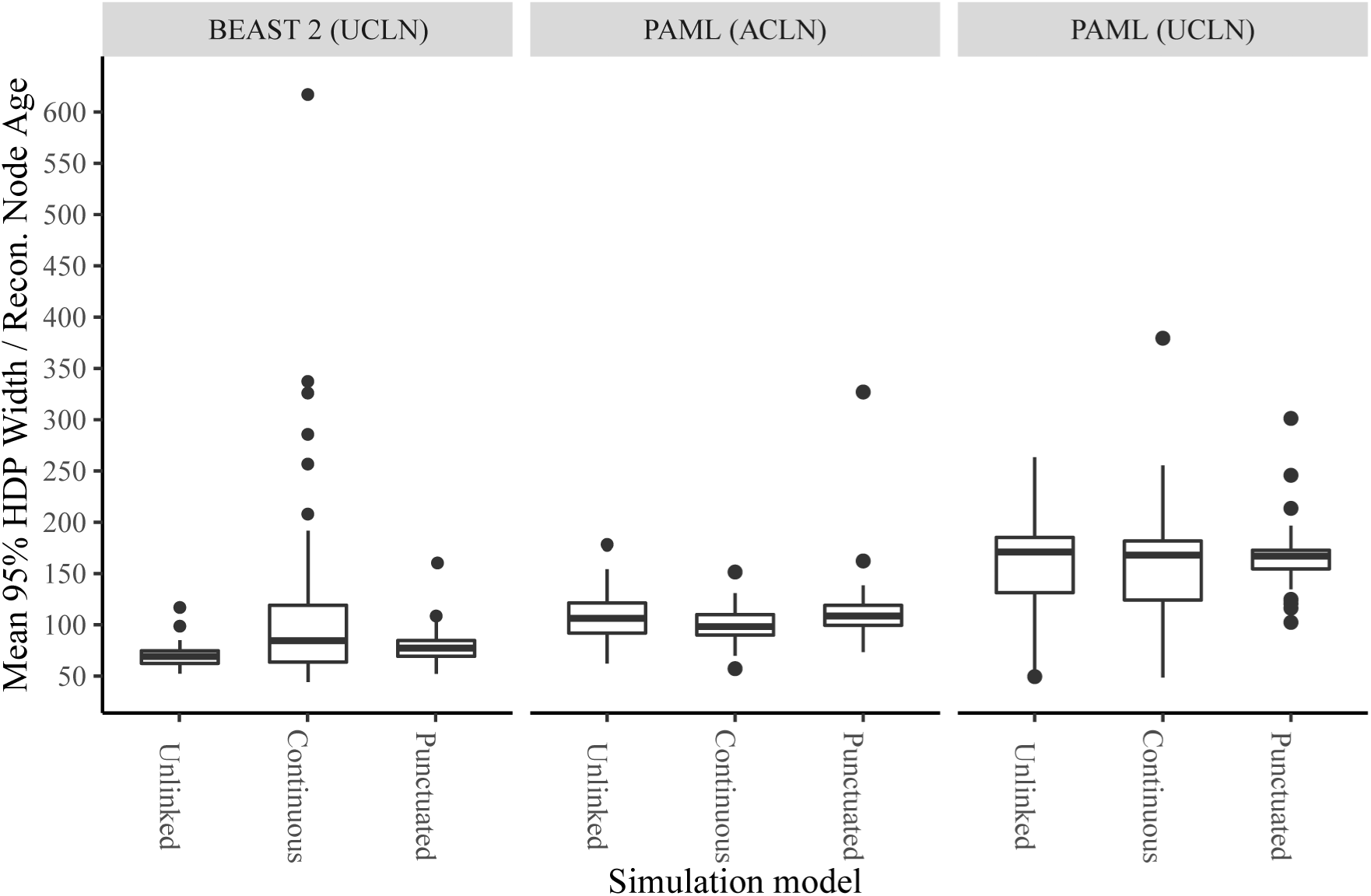
Uncertainty of trees reconstructed from phylogenies simulated under three models with different relationships between molecular evolution and speciation, in width as a percentage of reconstructed node age. Precision is measured by averaging the width of 95% highest posterior density intervals as a proportion of reconstructed node height across each tree, with a larger score indicating less precise estimates (HPD width). The three simulation models are Unlinked (instantaneous covariance of molecular rates and speciation rates = 0), Continuous (instantaneous covariance = 0.0044), and Punctuated (instantaneous covariance = 0, bursts of substitutions added at speciation events). Topologies and node times were reconstructed using three different analytical methods, the uncorrelated lognormal ‘relaxed clock’ (UCLN) rate prior in BEAST 2, the autocorrelated lognormal rate (ACLN) prior in PAML, and the UCLN in PAML.

Third, we calculated the uncertainty of node age reconstruction as reported by the method via the width of 95% highest posterior density intervals as a proportion of node age. Division by node age is required because the infinite-sites model of molecular evolution predicts that the relationship between posterior mean node age estimates and 95% credible intervals should approach a straight line with sufficient molecular data (75, 77). Using BEAST 2 (UCLN) (Figure 4), the average HPD width for the Unlinked trees had a width 60% of the reconstructed age of its associated node (43 – 29%). For the Continuous trees this was 108% (35 – 605%), and for the Punctuated trees it was 70% of node age (42 – 154%). Using PAML (ACLN), the average HPD width for Unlinked trees was 107.0% of node age (62.2 – 178.4%); for Continuous trees, 100.4% (57.2 – 151.5%); and for Punctuated trees, 112.4% (73.4 – 327.0%). Using PAML (UCLN), the average HPD width was greater than for PAML (ACLN) for all three simulation models, at 160% (49.5–263.4%) of node age for Unlinked trees; 158% (48.7–379.5%) for Continuous trees; and 165% (102.4–301.2%) for Punctuated trees.

**Figure 4.**
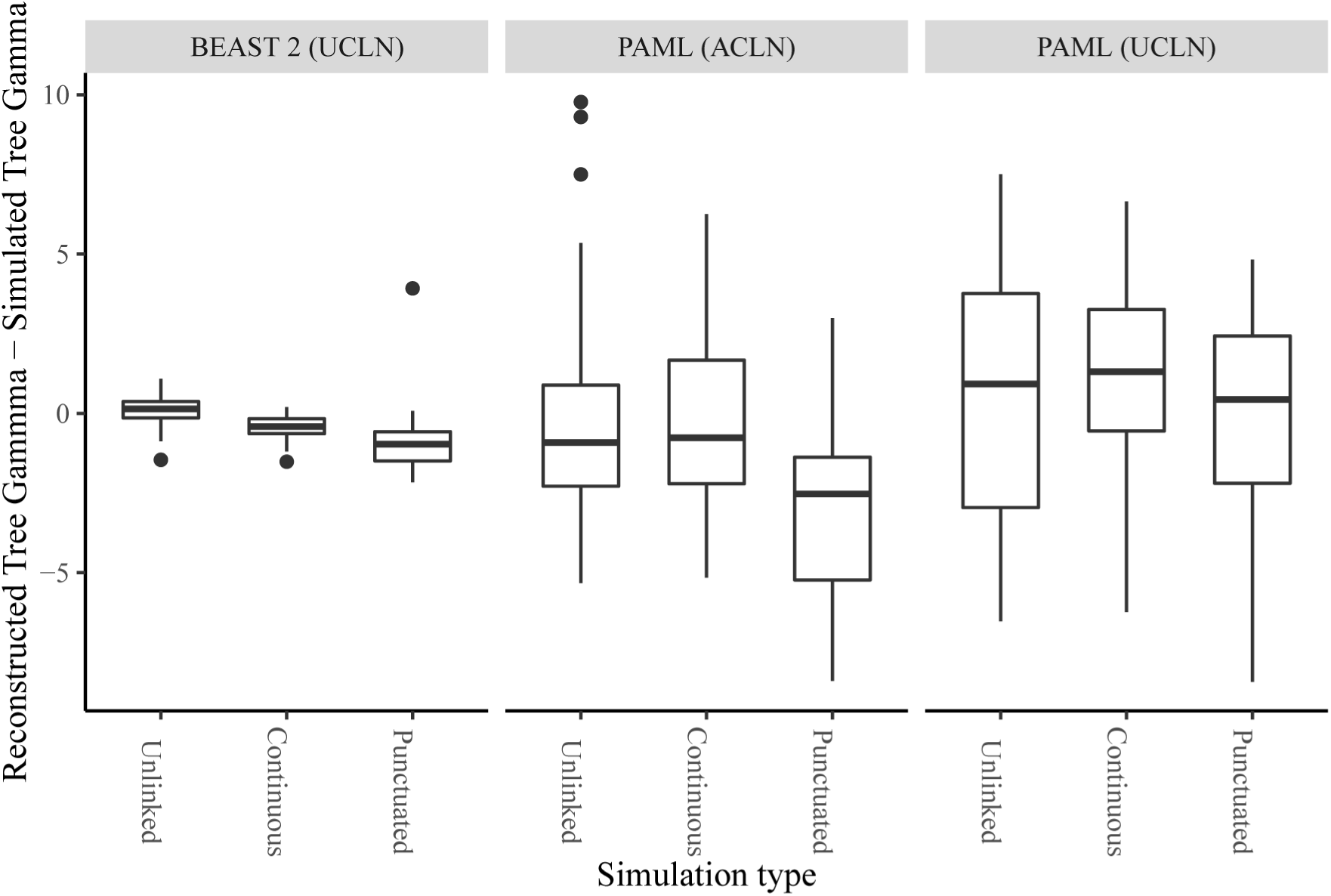
Difference in gamma statistic values between reconstructed and simulated trees, where simulated trees were generated under three models with different relationships between rates of molecular evolution and speciation rates. Negative scores indicate that reconstructed trees have nodes distributed more towards the root (‘tippier’) than the simulated tree. The three simulation models are Unlinked (instantaneous covariance of molecular rates and speciation rates = 0), Continuous (instantaneous covariance = 0.0044), and Punctuated (instantaneous covariance = 0, bursts of substitutions added at speciation events). Topologies and node times were reconstructed using three different methods, an uncorrelated lognormal (UCLN) rate prior in BEAST 2, an autocorrelated lognormal rate (ACLN) prior in PAML, and the UCLN in PAML.

### 2.4 Distribution of reconstructed vs simulated node ages

If the difference in gamma values between reconstructed and true trees is negative, the reconstructed tree has nodes distributed more towards the root, while if it is positive the reconstructed tree has nodes distributed more towards the tips than the true tree. Using BEAST 2 (UCLN), gamma values for the reconstructed Unlinked trees were distributed evenly around the gamma value of the simulated tree with a mean shift of +0.11 (-1.46 – + 1.09; Figure 5). This distribution was skewed negative (towards the root) relative to the simulated tree for Continuous trees with a mean of -0.44 (-1.52 – +0.20). This negative skew was more pronounced for the Punctuated trees at -0.94 (-2.16 – +3.92). Using PAML(ACLN), the relative change in gamma was on average negative across all three simulation models. For the Unlinked trees it was -0.28 (-5.32 – +9.77), while for the Continuous trees it was -0.76 (-5.15 – +6.25), and the Punctuated trees were again skewed negative with a mean of -2.87 (-8.40 – +2.99). In contrast when using PAML (UCLN), the relative change in gamma was on average positive for the Unlinked trees at +0.57 (-6.53 – +7.51) and for the Continuous trees at +1.07 (-6.24 – +6.65). It was slightly negative for the Punctuated trees at -0.16 (-8.43 – +4.83).

**Figure 5.**
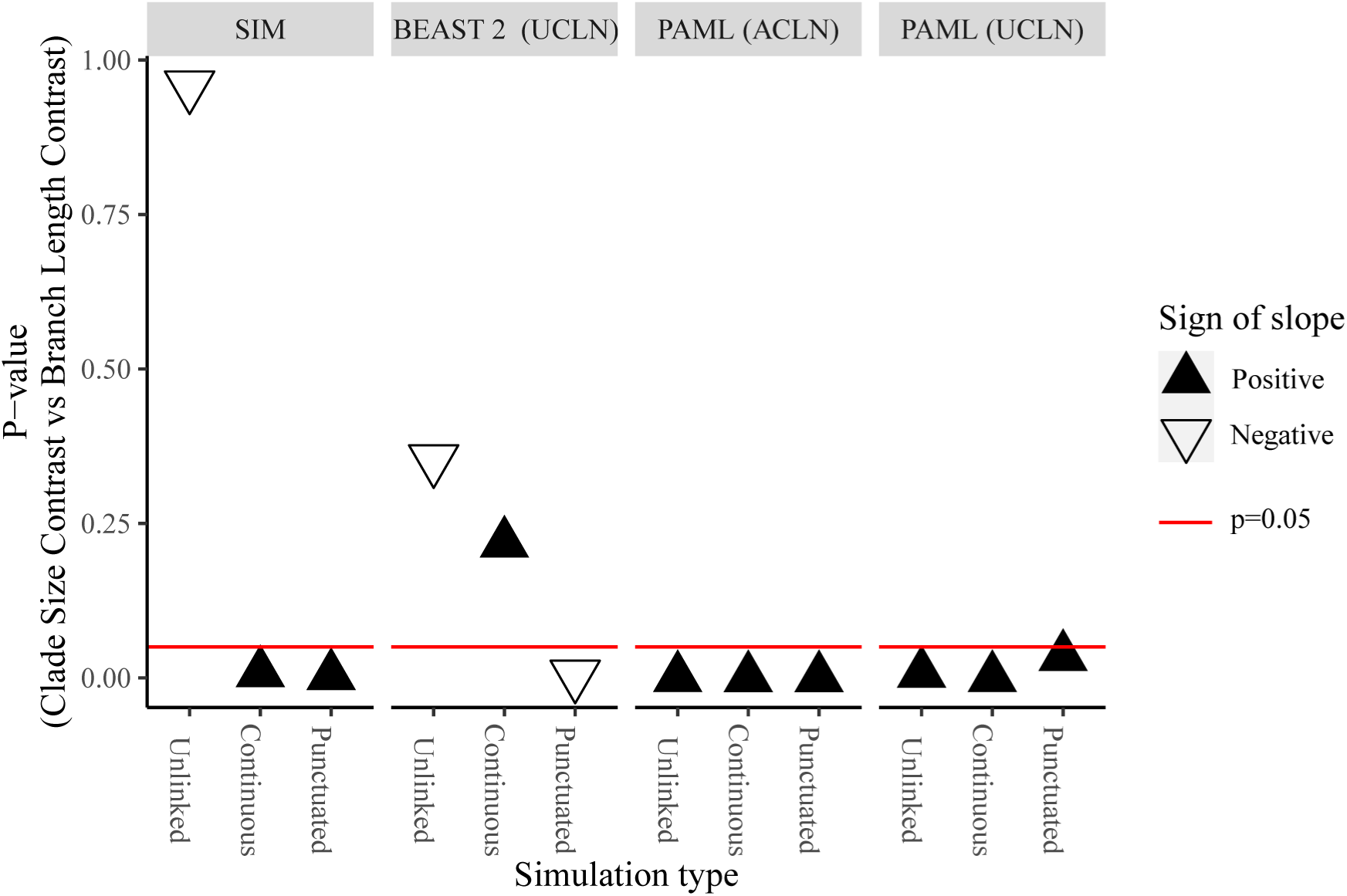
Slope and significance of mock sister pair analyses of clade size contrasts against contrasts in simulated branch lengths (SIM) or median branch length estimates reconstructed from simulated trees. Results are shown for branch lengths reconstructed using an uncorrelated lognormal (UCLN) rate prior in BEAST 2, an autocorrelated lognormal (ACLN) rate prior in PAML, or the UCLN in PAML. The three simulation models are Unlinked (instantaneous covariance of molecular rates and speciation rates = 0), Continuous (instantaneous covariance = 0.0044), and Punctuated (instantaneous covariance = 0, bursts of substitutions added at speciation events). Black triangles represent positive regression coefficients, while white inverted triangles represent negative coefficients. The red line indicates significance at the 0.05 level.

### 2.5 Detection of rate differences on reconstructed phylogenies

Sister pairs analysis have been used to detect an association between diversification rates and rates of molecular evolution in a wide range of datasets (6-8, 48, 88). In order to determine whether the simulated relationship between molecular rates and diversification rates could be detected in reconstructed trees, we regressed log clade size contrasts against contrasts in log median branch length estimates in the data set of sister pair analysis generated from reconstructed trees. We present the p-value and sign of slope for each regression (Figure 6). First, we asked whether the sister pairs analysis suggest a relationship between rates of molecular evolution and speciation rate in the simulated data (that is, in the true trees). As expected, Unlinked simulated trees show no significant relationship between clade size and branch length. Significant positive relationships are found for the Continuous and Punctuated simulations; note that this was a condition for selection of the Continuous data set (Supplementary Methods). Second, we asked whether the relationship between rates of molecular evolution and diversification rate could be detected in the reconstructed phylogenies. When reconstructed using BEAST 2 (UCLN), the Unlinked data set and the Continuous data set show no significant relationship, while the Punctuated data set gives a significant negative relationship, the opposite to what we expect. When reconstructed under PAML(ACLN) and PAML (UCLN), all three of the Unlinked, Continuous and Punctuated data sets have significant positive relationships between clade sizes and median branch length estimates.

**Figure 6.**
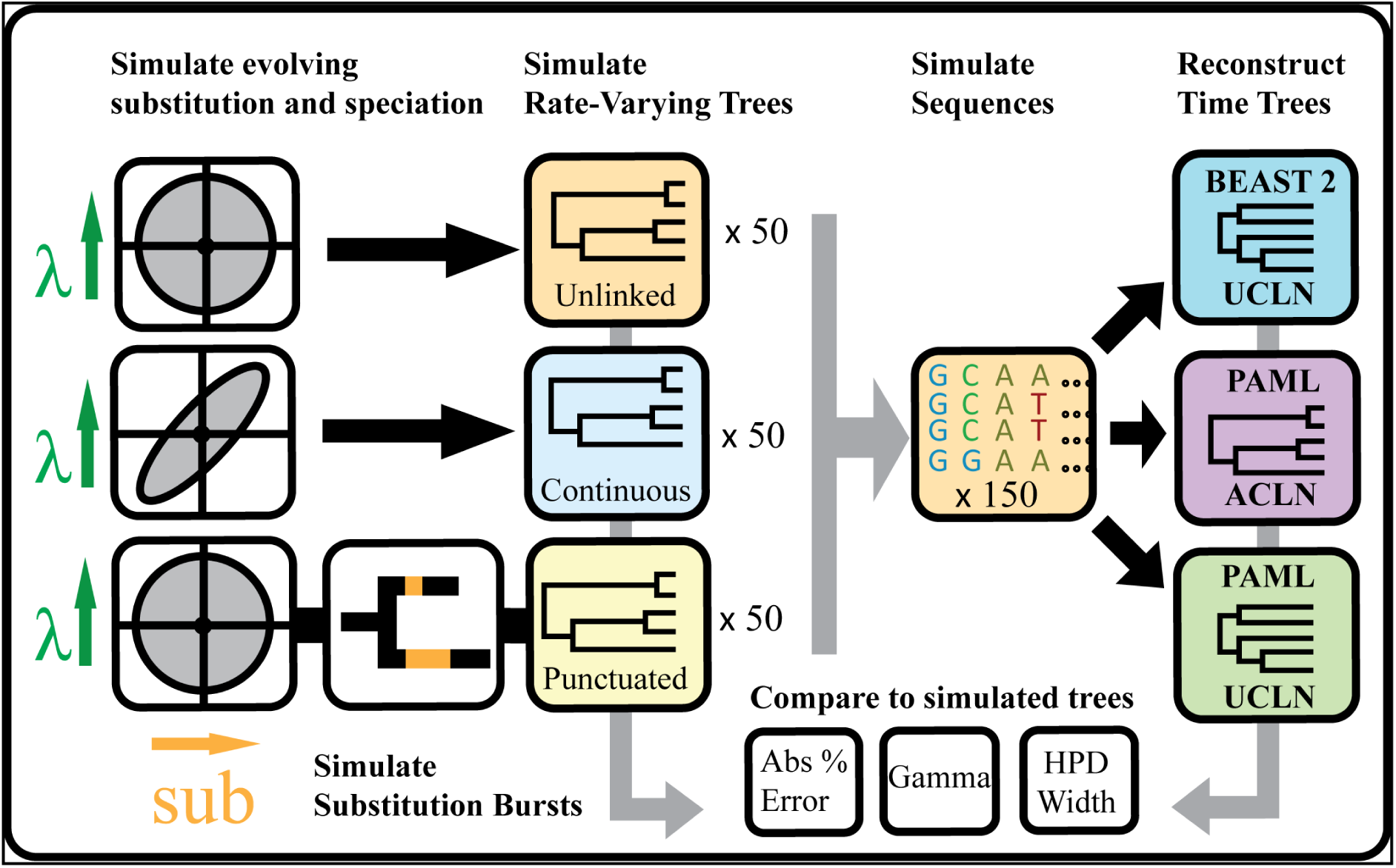
Flowchart of study design. Trees with branch lengths in units of time are simulated under three simulation conditions. For the Unlinked model, birth rates (*λ*)and substitution rates (sub) vary independently, while for the Continuous model they are correlated. For the Punctuated model, trees are simulated as for Unlinked, then some of the branch length is redistributed to form bursts of substitutions occurring upon speciation. Trees and branch lengths are then used to simulate molecular sequence alignments. Time trees are then reconstructed using three different methods with two rate priors: theuncorrelated lognormal in BEAST 2 (UCLN), the autocorrelated lognormal in PAML (ACLN), and the uncorrelated lognormal in PAML (UCLN). Reconstructed trees are then compared to the original, simulated trees to determine the error (Median absolute % error, MAPE), uncertainty (HPD width) and bias towards over- or under-estimation (Gamma).

## 3. Discussion

### 3.1 Dating errors associated with variation in speciation and substitution rate

The addition of rate-variable ‘relaxed clock’ models represents a significant extension to the sophistication of modern molecular dating methods (15). While there is a wide variety of rate-variable molecular dating methods, all of them to date rely on stochastic models of rate variation, such that rates of molecular evolution are expected to evolve randomly over the phylogeny. However, it is clear that real substitution rates do not always vary randomly. Instead, rates of molecular evolution can be influenced by species life history, niche and environment and macroevolutionary processes. It is important to examine whether the stochastic models of rate variation can adequately model realistic biological patterns of rate variation (3, 62, 65, 89). In the case of substitution rates, this means that we must test molecular dating methods using rate simulations that are empirically based and incorporate the effects of other biological processes. Here we have focused on testing the impact of widely reported correlations between molecular rates and diversification on the accuracy of molecular date reconstruction (5–8).

We used three models, Unlinked, Continuous and Punctuated, to reflect different possible links between substitution rates and diversification rates – either varying independently; covarying continuously, linked directly by common mechanisms or indirect through shared influences on rate variation; or linked by punctuated bursts of increased substitutions related to speciation events. Rather than choosing parameters arbitrarily, we parameterised our models using realistic values from the literature, or found plausible values by tuning the models to produce data sets that reproduce empirical findings (Supplementary Methods; 7, 79, 80). To make sure that we had realistic parameters for simulations, we based our parameter values on empirically derived values for birds. This approach produces more realistic simulations than the common approach choosing arbitrary values for convenience.

Although these values may not represent all taxa, they are still more likely to be a realistic representation than random values, and here are used only to explore whether molecular dating methods are accurate given simulated datasets that mimic, as much as possible, a real case study. Exploring the performance of methods on simulations modelled on other case studies would be desirable, if appropriate parameters were available. Modelling population genetic processes and multiple loci could also add real-world error sources such as gene tree discordance, coalescent error and incomplete lineage sorting, many of which would be affected by variation in substitution rates and branch times. Our framework focuses on the most common use cases for macroevolutionary studies, which generally involve analyses of few loci at broad taxonomic scales for which these problems may be less prevalent. However, this would be a useful path for future research.

Similarly, we use three examples of ‘relaxed clock’ molecular dating methods: BEAST 2 (using an uncorrelated rates model, UCLN) and PAML (using either the autocorrelated rates model, ACLN, or the UCLN). Clearly, these methods and models do not represent all possible approaches to inferring molecular dates using a rate-variable method, but they were chosen because they are widely used and differ in some key respects, and therefore present two useful test cases for the accuracy of “relaxed clock” methods in the face of systematic rate variation.

### 3.2 Unlinked model

The Unlinked model describes the error that can be expected from independent variation in speciation and molecular rates. Inference on trees simulated under this model produced median absolute errors in node age inference of about 12% on average using BEAST 2 (UCLN), and high values of 66% using the PAML (ACLN) and 77% using PAML (UCLN) (Figure 2). The largest errors were up to 30% for BEAST 2 (UCLN),while the worst instances for PAML (ACLN) had errors up to 192% and the worst MAPE for any individual tree was in PAML (UCLN) at over 500%. The Unlinked model produced lower errors using BEAST 2 (UCLN) than the other two models(Figure 2). Gamma differences for the Unlinked model are distributed evenly around zero using BEAST 2 (UCLN), so there is no overall tendency for nodes to be over- or under-estimated (Figure 4). The sources of error in the Unlinked model are most likely to be violation of the assumptions of rate priors and node time (tree) priors. Variation in speciation rates violates the assumptions of the birth-death node time priors used in both BEAST 2 and PAML, which assume universally constant rates of speciation and extinction (90). Speciation rate variation among lineages also induces more imbalanced trees on average (91), which can lead to bias and loss of precision in node age inferences (92). While the degree and scale on which speciation and extinction rates vary remains contested (93–96), there is increasing evidence that radiations and other forms of variation are a factor in the evolution of many taxa, so that errors of this magnitude may be widespread (97–100). Substitution rates that evolve in an autocorrelated fashion also violate the assumptions of the UCLN, which assumes that the evolutionary rate of each lineage is independent (20), and this may also induce errors in inference (30, 34).

The causes of the extremely high errors of the PAML(ACLN) and PAML (UCLN) phylogenetic inference for the Unlinked trees (and throughout the study for the other models) are unclear. Examination of the output trees suggests that the most extreme errors are in trees with shallower calibrations, and that the ages of large uncalibrated clades are sometimes overestimated severely. The inferences from PAML and BEAST 2 are not directly comparable due to numerous technical differences: for example, tree topology is fixed in PAML but estimated in BEAST 2; and BEAST 2 has prior distributions on speciation and extinction parameters, which are fixed in PAML. Nevertheless it is surprising that the performance of the ACLN model in PAML was worse than the UCLN model in BEAST 2 even though the underlying rate variation in these simulations is indeed autocorrelated. The fact that even higher errors and greater HPD widths are found when using PAML (UCLN) than using PAML (ACLN) (Figures 1, 3) suggests that the choice of rate prior can have some ameliorating influence, but the differences between the methods are not wholly due to the rates prior, because PAML (UCLN) still performs significantly worse than BEAST2 UCLN, indicating that there must be additional differences in methodology impacting the accuracy of inference. One possibility is that the sensitivity of the prior to model violation in PAML (ACLN) was enhanced by our use of the approximate likelihood function (101). This is a second-order Taylor approximation to the likelihood evaluated at the maximum likelihood estimates of the branch lengths and other parameters. As discussed by dos Reis and Yang (101), while the approximation is very accurate under normal circumstances, when the branch lengths proposed during Bayesian inference are very far from the maximum likelihood values the approximation becomes much less accurate. This can happen when the rate or node time (tree) priors are misspecified. PAML also requires some parameters of the prior to be specified exactly, such as the speciation and extinction rates, whereas in BEAST 2 these values are usually given distributions. This could mean that PAML is not as robust to violation of the assumption of constant diversification rates. Nevertheless, absolute error rates in other simulation studies have been less than 20% even with a deliberately misspecified rate prior and more limited calibrating information (30), and the high error rates may be indicating that patterns of rate variation likely to exist in real data may reveal weaknesses in methods that were not obvious from simulating simpler, stochastic patterns of rate variation.

**Figure 1.**
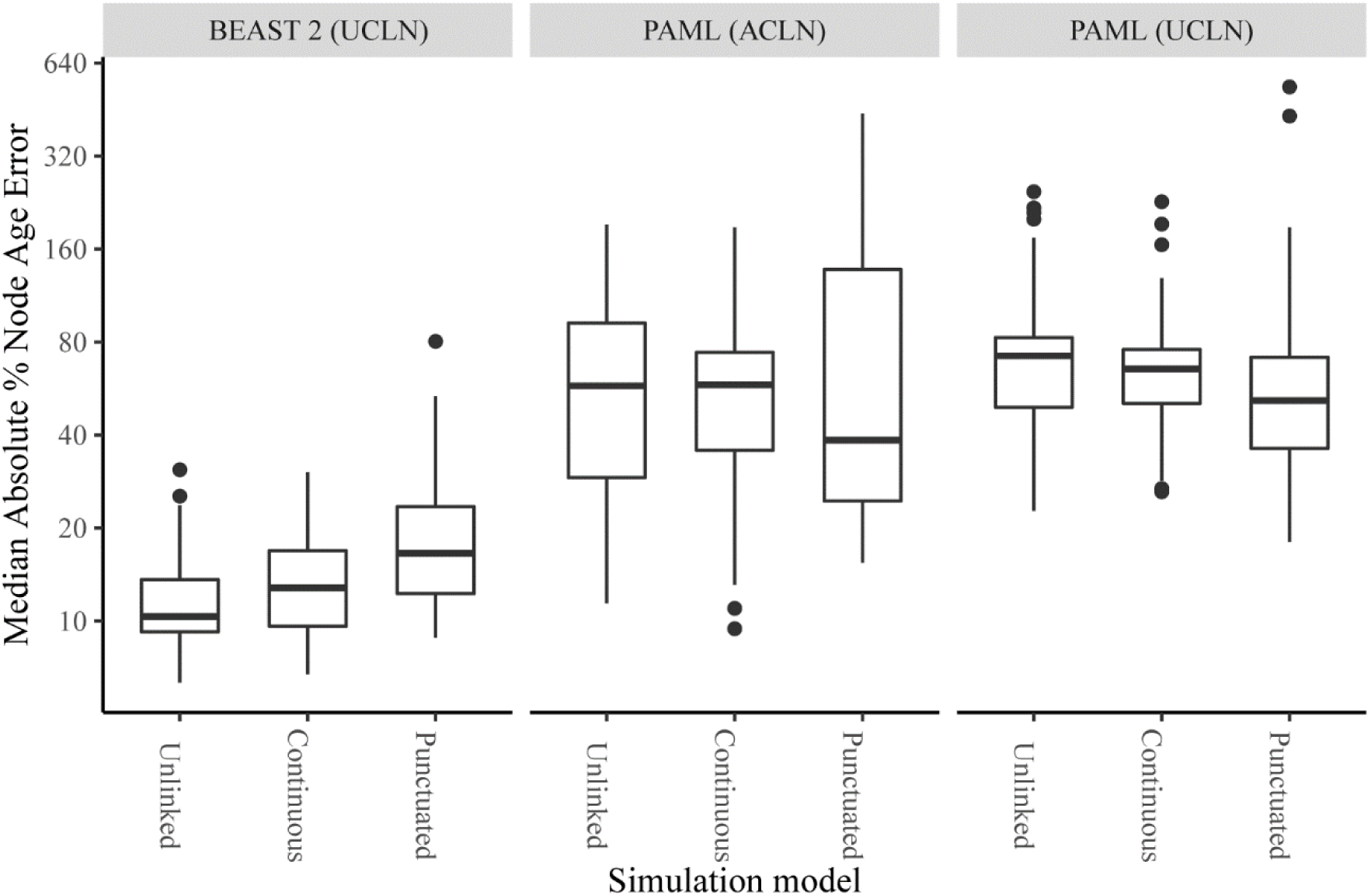
Median absolute percentage error of node times (on log scale) reconstructed from data sets simulated under three simulation types with different relationships between molecular evolution and speciation. The mean error is calculated only for branches whose bipartitions are shared between the true and inferred trees, and therefore ignores errors in topology which may occur in the BEAST 2 analyses. The three simulation models are Unlinked (instantaneous covariance of molecular rates and speciation rates = 0), Continuous (instantaneous covariance = 0.0044), and Punctuated (instantaneous covariance = 0, bursts of substitutions added at speciation events). Topologies and node times were reconstructed using three different analytical methods, with an uncorrelated lognormal ‘relaxed clock’ rate prior (UCLN) in BEAST 2, the autocorrelated lognormal rate prior (ACLN) in PAML, and the UCLN in PAML.

### 3.3 Continuous model

The Continuous model has substitution rates and speciation rates that coevolve continuously rather than varying independently. Notably, we chose the degree of covariance that reproduced two well-established empirical phenomena – the correlation of clade size and branch length contrasts detected in a sister pairs analysis by Lanfear et al. (7), and the widespread relationship between branch lengths and tree nodes reported by Pagel et al. (62). On average, this model produced absolute node time inference errors of 14% using BEAST 2 (UCLN), slightly higher than the Unlinked model, but the worst errors were up to 30% (Figure 2). Negatively-skewed gamma differences indicate that error generally leads to node ages being overestimated, pushing reconstructed nodes back towards the root (Figure 4). In addition to the error, which is the difference between reconstructed time estimates and the truth, we also examine how the reconstruction methods report their uncertainty about the truth through 95% highest posterior density intervals. Although not greatly different in terms of absolute error, the reported uncertainty of node time estimates was very high for this model using BEAST 2 (UCLN), with average HPD widths more than 107% of node height (Figure 5). This average is driven partly by some outlying trees with very high uncertainty, up to 600% of node height; the median is 75% of node height. Wide HPD intervals can be a sign of conflict between the prior and data, so it may be that covariance between diversification rates and substitution rates tends to produce patterns of branch lengths that are not well described by independent rate and time priors. The practical impact of this level of uncertainty could depend on how it is distributed within the tree and the nature of any downstream inferences; if the question at hand is the dating of major clades then high uncertainty in the age of tips may be tolerable, while the same level of uncertainty in internal branches would render the tree unusable. However, any macroevolutionary inference that relies on the distribution of nodes throughout the tree would likely be affected (59). Error rates for the Continuous simulations were similar to those for the Unlinked simulations using PAML (ACLN) (mean of 60.6%; Figure 2) and PAML (UCLN) (mean of 69.6%). Similar error rates suggest that the accuracy of PAML analysis, while having low precision under all simulation models, was not substantially affected by the degree of correlation between substitution rates and diversification rates.

### 3.4 Punctuated model

Levels of absolute error were highest under BEAST 2 (UCLN) when speciation events were associated with large punctuational bursts of molecular evolution (mean of 20.3%, up to 80.4%; Figure 2). This implies that, if real evolutionary processes were to commonly involve such bursts of substitutions associated with speciation, or produce a similar distribution of evolutionary histories and rate variation through some other mechanism, median node age estimates reconstructed under the UCLN rate prior would be generally incorrect. Error rates were also extreme for PAML (ACLN) (mean 91.4%, max 438.9%; Figure 2) and for PAML (UCLN) (mean 74.9%, max 535%). These rates were slightly higher on average than the rates for Unlinked and Continuous models, though also depending more on the specific tree. The finding of high error rates in the Punctuated model corroborates a recent analysis (72), which showed that greater proportions of branch length associated with speciation events lead to greater errors in the age of the root. Here, we show that the effect is strong enough to impact the median absolute error of all node ages, even when the crown node is calibrated. The very negative skew of Gamma away from the simulated value shows that these internal node ages are severely overestimated, leading to very distorted trees with long tips (Figure 4).

Despite having the highest degree of branch length error using BEAST 2 (UCLN), the Punctuated model had the lowest degree of topological error (a median of 2 bipartitions not correctly reconstructed; Figure 3), comparing to the Unlinked and Continuous models (median of 5 and 6 respectively, Figure 3). Intuitively, this could be because rapidly diversifying clades will normally have many short branches, making their internal relationships more uncertain. The Punctuated model adds more substitutions to every branch regardless of its duration in time, and so helps resolve these rapidly diversifying clades. The precise impact of topological error is difficult to interpret, since our investigation does not distinguish among topological errors at different timescales and we do not designate any particular nodes as important. But the practical upshot of systematically lower topological error in the simulation with the most severe node age errors using BEAST 2 (UCLN) is that these errors will be difficult to diagnose. Major topological errors are easier to diagnose by checking against classical taxonomies or other information, while little is usually known about the dates of uncalibrated nodes ahead of time.

Given the errors in inference of phylogenetic branch lengths and topology under these rate models, and the aforementioned difficulty in diagnosing these errors from inferred phylogenies, would we expect to be able to detect an association between diversification rates and substitution rates in these reconstructed phylogenies? We asked if the underlying relationship could be detected using a standard analytical approach with sister pairs (Figure 6). Although the association between speciation rates and substitution rates was evident in the simulated trees, tree reconstruction using PAML(ACLN), PAML (UCLN) or BEAST 2 (UCLN) disrupts the relationship between clade size and branch lengths and clade sizes displayed by the underlying simulated trees (Figure 4). In the BEAST 2 (UCLN) phylogenies the relationship between speciation and molecular rates is inverted, giving a negative correlation between clade sizes and branch lengths. This could be related to the fact that lineages with many speciation events have higher variance in branch lengths in the BEAST 2 (UCLN) reconstruction (102) which means their total length could be more strongly constrained by the node time (tree) prior. Conversely, for the PAML (ACLN) and PAML (UCLN) reconstructed trees, we found significant positive relationships between inferred branch lengths and clade size for all three models, even the Unlinked model where speciation rates and substitution rates are not positively associated. A possible cause of this inference of a positive association between diversification rates and substitution rate, even in the absence of an underlying association between the two, is the node-density effect, which results from systematic underestimation of branch lengths with high uncertainty (103). The influence of the node density effect should be reduced by our procedure of selecting an even number of samples in each sister clade, but may not be eliminated by it due to the extremely high error rates in the PAML (ACLN) or PAML (UCLN) reconstructions. The node-density effect may not be observed in our BEAST 2 (UCLN) because the aforementioned artefactual negative correlation overcomes any positive correlation due to node density. However the reasons why this does not occur using PAML (UCLN) are unclear and cannot be distinguished using our data. The failure to recover the empirical correlation from branch lengths reconstructed as part of a Bayesian dating analysis suggests that this is not a useful way to test for the presence of interactions between substitution rates and speciation rates. For some data sets it may be practical to check for such correlations by reconstructing branch lengths in units of genetic divergence using a fast branch length reconstruction method that does not impose a model of timing or rate variation (e.g. 85, 104, 105, 106). However, even if such correlations can be diagnosed, there is currently no available method for correcting any associated bias. This should be an area of active research.

The true role of punctuational bursts or coevolution of speciation and substitution rates in macroevolution is currently a matter of debate. The primary empirical evidence is the analysis of a large selection of empirical phylogenies in Webster et al. (63) and Pagel et al. (62). These studies have found that the length of root-to-tip paths in the phylogenies in substitutions per site is frequently higher when there are more nodes on the path. The relationship remains even after removing the node-density effect that can generate spurious correlations between branch lengths and node densities across clades (103, 107). Regardless of the mechanisms, we have shown that if there were punctuated busts of substitutions associated with speciation events, we could expect it to impact the accuracy of molecular date estimates, since any mechanism that leads to similar patterns of rate variation and apparent correlation between branch lengths and speciation events would produce similar effects.

### 3.5 Molecular dating methodology

Our study contrasts two common reconstruction methods, rather than varying aspects of data choice. Many aspects of data choice and analytical method could exacerbate or reduce the errors found in this analysis. For example, our alignments are shorter than many used in actual Bayesian phylogenetic analyses but also likely more informative due to being simulated without invariant sites or other more complex patterns of substitution. Shorter or less informative alignments could lead to greater or more widespread error. Perhaps the most important element that we did not explore was the influence of calibrations. Calibration choice and positioning, as well as the shape of the calibrating distribution used, is often the most influential factor influencing divergence time reconstruction (e.g. 32, 108) and is the source of many disagreements in published molecular dating studies, including high-profile dating studies in birds (109–111) mammals (112, 113), and insects (114–116). Simulation studies have suggested that calibrations can rescue analyses strongly affected by rate prior misspecification (30), although others have found that sufficiently misspecified rate priors cannot be rescued by more calibrations (34). The largest and most data-rich molecular dating studies may have many more calibrations than we implement (e.g. 139, 117). However, our trees are smaller than in these studies, and, unlike empirical studies, our calibrations are guaranteed to be correctly placed, have hard constraints, and are centered on the true value. We also ensure the root is calibrated, which has been shown to have the largest effect on accuracy in several studies (30, 34, 118). We therefore believe that our method overall represents a balance between realism and conservatism in determining the amount of calibrating information available to inform divergence time estimates. However, a usefl extension of the analyses undertaken in this study would be investigation of the effects of both alignment length and calibration numbers and positioning.

## 4. Conclusions

Our results here show the value of testing molecular phylogenetic and dating methods against complex simulation models informed by empirically determined patterns of rate variation. We have demonstrated that commonly applied Bayesian divergence time estimation methods can experience average errors of 12% in node dates under a model in which speciation and molecular rates vary independently, 14% when substitution rates are linked to speciation rates and 20% under a punctuational model when analysed using an uncorrelated rates prior in BEAST (UCLN). Errors were much higher under all models when trees were analysed with an autocorrelated rate prior in PAML (ACLN), and still higher with the uncorrelated lognormal rate prior in PAML (UCLN). As demonstrated by the negative skewness of gamma statistics, for trees with links between speciation and molecular evolution these errors could lead to systematic overestimation of node ages. Regardless of the mechanism generating the association between speciation rates and substitution rates, we show that the potential for divergence time estimation error associated with known empirical relationships between molecular evolution (manifest in substitution rate) and macroevolution (manifest in the topology and branch lengths of phylogenetic trees) can be significant and should be taken into account. Future work should see a broader range of sophisticated divergence time estimation methods be tested against a wider variety of more empirically realistic simulated data, including geographic and environmental biases, concerted changes in population size, and directional changes in life history characteristics. Ultimately the goal should be to produce new models that relax the assumption of independence between priors on divergence times and priors on substitution rates. These will lead to greater confidence in applications that depend on robust estimates from molecular dating procedures.

## 5. Methods

### 5.1 Simulation design

We refer throughout to simulated phylogenies, which are known without error, as the ‘simulated trees’, and to the phylogenies inferred from analyzing the sequences evolved along each of these simulated phylogenies as the ‘reconstructed trees’. Comparison of the reconstructed trees to the simulated trees allows us to evaluate the degree to which phylogenetic inference methods correctly infer the known evolutionary history of the simulated data.

Our overall design is summarised in Figure 1. For our simulation study, we first generate phylogenetic trees in units of absolute time with evolving rates of speciation and molecular evolution. We then simulate DNA sequences along these phylogenies. Finally, we infer dated trees from the alignment of these sequences using standard methods in Bayesian phylogenetics and compare the inferred tree to the simulated tree from the known evolutionary history from each simulation.

We simulate sets of 50 phylogenetic trees and nucleotide sequence alignments under each of three different models describing the association between speciation rates and rates of molecular evolution: one “Unlinked” model in which molecular and diversification rates each vary independently, as well as two alternative mechanisms for an association between rates of molecular evolution and speciation rates (“Punctuated” and “Continuous”: see Table 1).

We do not consider models in which molecular rates affect rates of extinction, so in our simulations, extinction rates remain constant over the tree. For all models, we first simulate the phylogenetic tree in continuous time, together with speciation and substitution rates that vary along the tree. From this, we derive branch lengths in expected genetic divergence (substitutions/site). Finally, we simulate the evolution of nucleotide sequences along these phylogenies.

In the Unlinked model, we simulate continuous variation in both substitution rate and speciation rate through time, but neither affects the other. Substitution and speciation rates evolve independently by Brownian motion, where each lineage continuously adds a separate random increment to its substitution rate and speciation rate in each of many small time intervals. The Unlinked model serves as a baseline to determine the level of uncertainty in molecular dates caused by the random variation in substitution and diversification rate when there is no mechanistic link between the two. In the Continuous model, speciation rates and substitution rates coevolve with each other continuously over time. The coevolution is simulated as a correlated Brownian motion, where the random increments in speciation and molecular rates added over time are dependent on one another. Consequently, substitution rate and speciation rate tend to increase or decrease together. For this model, we also ensured realism simulated many data sets with different covariance parameters, and selected a set of 50 trees that reproduced the empirical correlation of (7) between clade size and branch length (Supplementary Methods S1.1). In the Punctuated model, the genetic divergence along a branch is the sum of a gradual accumulation of substitutions at a rate that evolves as in the Unlinked model, and a burst component at each speciation event. A speciation event occurs each time a new lineage is born, whether any of its descendants are observed in the present or not. The amount of additional branch length due to the burst is independently and randomly drawn for each speciation event. For this model, we condition our simulations to reproduce the relationship observed in (62). There it was observed that the total number of substitutions occurring along root-to-tip paths in their sample of trees correlates with the number of nodes along the path. Although this relationship could also be produced by a mechanism similar to the Continuous model (119–121), we model the mechanism proposed in (62) that the node-associated substitutions are produced in bursts associated with speciation events.

We derived plausible speciation, extinction and substitution rates for our simulations using the avian phylogeny in (82). We do not intend to make any inferences about the actual evolution of birds or the accuracy of inferred divergence times. Birds are chosen only because the relationships among traits, molecular rates and diversification rates are well studied, so that we can use these to give us an empirical ground for choosing parameter values. Branch-specific diversification rates have been estimated for many of the lineages in this tree in (80), while substitution rates have been inferred in (79) assuming covariation of molecular rates and life history. Speciation rate and molecular rate values taken from avian data have been previously used to inform realistic simulations on ecological assemblies for the purpose of estimating the effect of dating error on phylogenetic diversity estimates (122), and we make use of similar procedures here. A relationship between molecular rates and net diversification rate has also been established for birds (7, 8). We use this fact to select our Continuous data set by simulating multiple data sets and varying the covariance parameter until the relationship appears. Full details of how we used these studies to determine realistic parameterisations for our simulations are available as Supplementary Methods S1.1.

### 5.2 Simulation trees and alignments

To simulate the phylogeny, we adopt a forward approach, starting at the root of the phylogeny and moving forward in time, as lineages divide into two daughter lineages, go extinct, and evolve in speciation and substitution rate as a continuous time Markov process. In each step, we draw the time until the next event from an exponential distribution with rate equal to the sum of the rates of all events in all lineages. Then, for every lineage *l* that is present at time *t*, one of the following three events may take place:

(i) The lineage gives rise to two daughter lineages with rate *λ*(*l,t*), the current speciation rate of lineage *l* at time *t.* Both daughter lineages initially retain the same speciation rate and trait values as their parent lineages.
(ii) The lineage becomes extinct with rate *μ*, which is constant over the tree.
(iii) The lineage updates its speciation rate and substitution rate with rate *q*, which is constant throughout the tree. The update process adds an increment to the current speciation rate, which is a random draw from a bivariate normal distribution with mean 0 and an instantaneous variance-covariance matrix *Σ* multiplied by the time since last jump. Until one of these events happens, the lineage continues with the same speciation and molecular rates. High values of *q* make for longer simulations but produce a better approximation to Brownian motion. We found that *q* had no influence on observed speciation or substitution rate variance or average tree age for *q >* 10. For our final simulations, we selected *q* = 50 to allow a significant margin of safety.

Specifically, the instantaneous variance-covariance matrix takes the form:

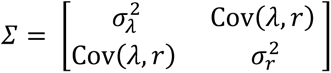

where 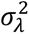 describes how fast the speciation rate evolves and 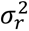 describes how fast the substitution rate evolves. Cov(*λ,r*) = *ρσ*_λ_*σ_r_*, where *ρ* is the correlation coefficient between speciation rate and substitution rate, so *ρ* = 0 in the Unlinked Model and the Punctuated model. We employ the Generalised Sampling Approach (123) to ensure a correct distribution of trees with a given number of tips, which is set to 75, the approximate mean size of the avian clades taken from (80). Details of the approach is in Supplementary Methods S1.2.

In the Punctuated model, we simulate an initial tree as per the Unlinked sample, but without removing extinct taxa. We generate a burst of substitutions following each speciation event. We generate Punctuated trees to match the formula found by (62), in which 16 ± 5.4% of the tree length (sum of all branch lengths) results from node-associated bursts (Supplementary Methods S1.3).

Once we have simulated a tree, we simulate the evolution of nucleotide sequences along the tree, given that branch lengths in the tree represent the expected number of substitutions per site along the branch. The branch lengths in this tree represent the expected number of substitutions per site along the branch. We simulate alignments of 2000 bases, which is roughly equivalent to a 6000-base coding alignment, since most substitutions in a coding alignment occur in the third codon sites. A 6000-base coding alignment is about the size of the concatenated coding sequences of a mitochondrial genome. Previous literature reviews have shown this to be a common length for phylogenetic data sets in published ecological studies, and a tractable length for large Bayesian simulation studies (122). We parameterize our sequence simulations using realistic among-site rate variation values from birds (Supplementary Methods S1.4).

### 5.3 Calibrations

In addition to the trees and sequence alignments, we need calibrations to provide information on absolute time to allow substitution rates to be identifiable from absolute ages of nodes. For simulated trees of similar size to ours, diminishing accuracy returns for adding new calibrations are achieved between 1 and 5 calibrations, especially if one is at the root (30). Since our calibration dates are known with more accuracy than those in a real analysis, and we remove the possibility of incorrectly assigned calibrations, we choose to assign two calibrations per tree. For each tree, we choose calibrations at the root node and at one randomly chosen internal node. To ensure that the calibration is randomly chosen across the nodes of the simulated trees, we first decide the level of the calibrated node as a random integer between 1 and the largest number of nodes along any root-to-tip path in the tree. Then we randomly pick the calibrated node from all the nodes at that level.

For analyses in both BEAST 2 and PAML, we apply a hard-bounded uniform prior for the age of each calibrated node. Both calibrations have bounds at the true age ± 15%. Note that this is a conservative test of molecular dating because in the simulated datasets we have accurate knowledge of the true date of the speciation event. In reality, calibrations may be misplaced or incorrectly dated, and may only provide maximum or minimum bounds.

### 5.4 Phylogenetic inference

We generate a set of 50 trees under each of the Unlinked, Continuous and Punctuated simulation types. After sequence simulation, this gives us 50 sequence alignments for each of the three models.We infer divergence times from the simulated sequences using common Bayesian molecular dating procedures. First, we generate posterior distributions of tree topologies and divergence times using three common molecular dating methods: the uncorrelated lognormal rate prior (UCLN), as implemented in the software package ‘BEAST 2’ (BEAST 2; 83), in which rates for each branch are independent draws from a common lognormal distribution;the autocorrelated lognormal rate prior (ACLN), implemented in the ‘mcmctree*’* program in the PAML software suite (85), in which the rate of each branch is lognormally distributed with a mean equal to the rate of its parent branch; and the UCLN as implemented in ‘mcmctree’ in PAML. To speed analysis when using PAML, we used the approximate likelihood algorithm implemented in mcmctree. The PAML analyses also differ from the BEAST 2 analyses in that some model parameters are provided as fixed numbers rather than distributions, and that PAML requires a fixed tree topology. In this case the fixed topology was the topology of the true simulated tree. Therefore only BEAST 2 (UCLN) incorporates the possibility of topological inference errors. However, the three simulation models can be compared within each reconstruction method. Details of the inference method are available as Supplementary Methods (S1.5).

### 5.5 Error in reconstructed trees

For each reconstructed tree, we calculated accuracy in the estimation of node times using the median absolute percentage error (MAPE) of inferred node times for clades that are present in both true and reconstructed trees. The median was chosen rather than the mean to reduce the impact of extremely high values for the estimates of shallow nodes. Since these are only calculated for correctly reconstructed branches and do not capture errors in reconstructing topology, we also calculate the Robinson-Foulds distance or number of non-shared bipartitions, which only reflects topological error (87).

Model misspecification in Bayesian molecular dating methods can sometimes be associated with higher uncertainty in the reconstructed node ages, as measured by the width of Bayesian credible intervals. This is undesirable because the reconstructed history is less useful as an explanatory tool and less informative in downstream analyses, but it can also be a simple way to diagnose problems with the specified model. We examined the uncertainty with which node ages were reconstructed by the two Bayesian molecular dating methods. We scored uncertainty by averaging the width of the reconstructed 95% highest posterior density intervals as a proportion of reconstructed node height across each tree (HPD width), with a larger score indicating less precise estimates.

### 5.6 Distribution of reconstructed vs simulated node ages

We examine the direction of error by calculating the gamma statistic for each tree and comparing to the true tree (124). The gamma statistic is a description of how the times between branching events in a tree whose tips are all extant taxa change as they become closer to the present. It summarises information relating to the pattern of diversification, following a standard normal distribution for a tree with a constant rate of speciation and no extinction. Trees with higher values have a relatively greater proportion of their length in older branches, while those with lower values have relatively more of their length in younger branches. The statistic therefore tells us whether the ages of internal nodes within each tree tend to be older or younger on average than the true times while accounting for the different scale of this bias for older and younger nodes.

### 5.7 Efficacy of sister pair correlation on reconstructed branch lengths

We checked to see if we can use the reconstructed branch lengths, which are the median of posterior branch lengths calculated by BEAST 2 (UCLN), PAML (ACLN) or PAML (UCLN), to detect the correlation between branch lengths and clade sizes found by sister pair analysis on the simulated trees (see Supplemental Methods available as online supplementary information). This could be used as a rapid post-hoc check to see whether published dated trees could be affected by reconstruction error that is associated with substitution and speciation rate covariation. Data sets for the sister pair analysis are generated from the 50 simulated trees under each of the three simulation models, where each tree gives a pair of sister clades and the sister clades are the two clades that split at the root. So, under each simulation model, we have a data set of 50 sister pairs. We then regress contrasts in log clade sizes against contrasts in the log of the median branch length estimates from BEAST 2 (UCLN), PAML (ACLN), and PAML (UCLN).

## Supporting information

Supplemental methods and results

## Abbreviations

UCLN: Uncorrelated Lognormal
ACLN: Autocorrelated Lognormal
HPD: Highest Posterior Density
MAPE: Mean Absolute Percentage Error

## Declarations

### Ethics approval

Not applicable.

### Consent for publication

Not applicable.

### Availability of data and materials

All data generated during this study are included in this published article as Supplementary Information. The datasets generated are also available online at https://github.com/amritchie/Sim_Evolving_Rates_Molecular_Dates.

## Funding

This work was supported by the Australian Research Council grant DP190101916.

### Authors’ contributions

The study was conceived by L.B. A.M.R. wrote simulation code and conducted the analysis.

X.H. reviewed simulation code and assisted in the design of the study. A.R. and L.B. wrote the manuscript. All authors reviewed the manuscript.

## Acknowledgements

We would like to extend our gratitude to Marcel Cardillo and Alex Skeels for helpful conversations in the preparation of this manuscript.

