## Supplemental methods and results for "Investigating the reliability of molecular estimates of evolutionary time when substitution rates and speciation rates vary"

SUPPLEMENTARY INFORMATION

Table of Contents:

**2:6 Supplementary Methods**

S1.1 Parameterising realistic rate variation

S1.2 Generalised Sampling Approach for sampling trees

S1.3 Simulating trees with punctuational bursts of substitutions

S1.4 Sequence simulation

S1.5 Details of molecular dating analyses

**7 Table S1 - Simulations conducted**

**8 Figure S1 – Results for alternative node age error metric (Branch Score Distance)**

Additional Supplementary Files:

**simulated_trees:** Simulated data sets of 50 phylogenetic trees in Newick format.

**simulated_alignments:** FASTA files containing nucleotide alignments simulated along the correspondingly numbered trees.

**beast_reconstructed_trees:** Trees reconstructed from simulated alignments under the Uncorrelated Lognormal Relaxed Clock (UCLN) in BEAST 2, with XML files included.

**paml_reconstructed_trees:** Trees reconstructed from simulated trees and aligments under the Autocorrelated Lognormal Relaxed Clock (ACLN) in mcmctree

**code_control_files:** R code for simulating sampled trees with covarying speciation and substitution rates (bdforward.R), and example control files for PAML (Unlinked_1.ctl, Unlinked_1_run2.ctl).

All simulation and analysis files are also available online at https://github.com/amritchie.

*S1.1 Parameterising realistic rate variation*

We estimated the rate of evolution of the speciation rate, $\sigma_\lambda^2$, from the bird tree of Jetz et al. (2012) using phylogenetically independent contrasts for speciation rates at the tips within each of the major bird clades analysed by Maliet et al. (2019). The phylogenetically independent contrasts are calculated for each pair of sister tips within each clade using the "pic" function implemented in the R package "ape" (Paradis and Schliep 2019). We calculated the variance within each clade as the average of these contrasts, then averaged these results across clades to produce our final variance parameter for speciation rates. For the constant extinction rate, we take the average of the values estimated per clade in Maliet et al. (2019), equal to 0.0046. Applying the same method we used to estimate $\sigma_\lambda^2$, we estimated the rate of evolution of substitution rates, $\sigma_r^2$, using phylogenetically independent contrasts of avian substitution rates estimated by Nabholz et al. (2016).

We do not have an empirical estimate of the covariance of speciation and molecular rates for use in the Continuous scenario. Empirically derived correlation coefficients from studies such as Lanfear et al. (2010) refer to the correlation of clade sizes with branch lengths in a set of pairs of sister clades, rather than the instantaneous covariance of molecular and speciation rates. They are also likely to be substantially weaker than the true underlying correlations due to information loss from taxon sampling and phylogenetic uncertainty. Nevertheless, these studies have shown that an association between diversification rate and rates of molecular evolution is significant and widespread (Bromham et al. 2015; Eo and DeWoody 2010 ; Iglesias-Carrasco et al. 2019). Therefore, to parameterise the correlated rates in our Continuous scenario, we choose a covariance that is able to reproduce this observed empirical relationship. This allows us to generate simulations that resemble the relationships observed in real datasets.

To select an appropriate value, we tested covariance values of 0.0011, 0.0022, and 0.0044, yielding theoretical correlations of 0.2, 0.4 and 0.8 (Table S1). We used 0.2 as the lowest correlation coefficient, because this was the correlation coefficient between clade sizes and branch lengths in Lanfear et al. (2010). The higher values are 0.2 doubled and quadrupled. We then simulated 10 mock sister pair data sets under each covariance value. In each data set, we simulated 50 trees and each of these trees gives one sister pair, where the sisters are the two lineages that split at the root of the tree. We then regressed the contrast in clade sizes of the sister pairs in each tree against the contrast in the average root-to-tip branch lengths in substitutions/site through the origin. All contrasts were standardised by the square root of the age of the pair (Garland et al. 1992). Of the three test covariances, only the data set simulated with a covariance of 0.0044 (theoretical correlation of 0.8) produced significant clade size vs. branch length contrasts in a majority of its 10 mock sister pair data sets (total of 9 out of 10). As an additional check on realistic patterns of rate variation in our simulated data, we used the test of Webster et al. (2003) to look for evidence of correlations between substitutions and speciation events on root-to-tip paths in individual trees. Of the total 500 trees across the 10 data sets simulated using the covariance of 0.0044, 176 (35.2%) showed a significant relationship between substitutions and speciation using this test. This is close to the 35% of trees with significant relationships found in the survey of Pagel et al. (2006). So, we found that the same covariance value under the Continuous simulation model can reproduce both the observed association between substitution rate and diversification rate (clade size) and the observed association between path length and number of nodes from root to tip. This finding is consistent with the argument by some authors that the link between substitution and speciation can be equally explained by a gradual mechanism and a punctuational mechanism (Pennell et al. 2014; Pennell et al. 2014; Rabosky 2012). For the Continuous simulation model, we selected a set of 50 trees at random from among the 10 sets simulated with a covariance of 0.0044 and showing a significant empirical correlation using the sister pairs test, for comparison with the 50-tree Unlinked and Punctuated data sets. 20 of these trees showed a positive correlation using the test of (Webster et al. 2003). As a result, we have $\sigma_\lambda^2$ = 0.015, $\sigma_r^2$ = 0.0044, and $\text{Cov}(\lambda, r) = 0.0021$ in the Continuous scenario and $\text{Cov}(\lambda, r) = 0$ in the Unlinked scenario and the Punctuated scenario.

*S1.2 Generalised Sampling Approach for sampling trees*

We employ the generalised sampling approach (GSA) to sample trees conditioned on the number *n* of extant taxa (Hartmann et al. 2010). *n* was set to 75, the approximate mean size of the avian clades taken from (Maliet et al. 2019). It does not suffice to stop the birth-death process when the desired number of taxa is achieved, because the sample will then not include trees that could have arrived at this number through extinction. Therefore, it is necessary to evolve trees forward to a number of taxa *m* that is greater than *n* and is unlikely to reduce to *n* due to extinction if simulation continues. Then, we simulate a large number of trees stopping at *m* taxa and sample each tree with a probability proportional to the amount of time the tree had *n* living taxa during the simulation. The tree is then cut at a time point chosen uniformly from all the time points during the simulation when the tree had *n* living taxa. Based on experimentation, we used values of *m* = 100 for *n* = 75.

*S1.3 Simulating trees with punctuational bursts of substitutions*

We start with branch lengths we generated in our tree simulations, representing the gradual component of the tree length $G$. We take the mean tree length as $T = \frac{G}{1-0.16}$, and draw the total branch length from a uniform distribution between $T \pm 0.054T$. The added length $T - G$ is distributed evenly among branches, including on extinct lineages, where the proportion on each branch follows a symmetric Dirichlet distribution. Lastly, we rescale all branches so that the final tree length is unchanged after adding bursts, and remove extinct taxa. This procedure results in some substitutions being lost as they are redistributed to extinct taxa, but this loss is not large because of the low and constant extinction rate. The average and standard deviation of tree lengths of the resulting Punctuated trees is $3.55 \pm 1.3$ compared to $3.61 \pm 1.2$ for the Unlinked trees and $3.84 \pm 0.96$ for the Continuous trees. The effect of this procedure is a redistribution of substitutions from longer to shorter branches in the proportion suggested by Pagel et al. (2006).

*S1.4 Sequence simulation*

Sequences of 2000 bases are simulated in R using the simSeq function in the R package 'phangorn' (Schliep et al., 2011). To simulate variation in substitution rates across sites, sites are first partitioned into rate categories as described by a discrete Gamma distribution with a shape parameter of 20, leading to an approximately 30-fold difference in rates across sites as is known to exist in avian mitochondrial control regions (Barker et al., 2012). These site rates are multiplied by the average branch rates to produce the final rate for each site partition. Ancestral nucleotide frequencies are drawn from a symmetric Dirichlet distribution, and sequences are then simulated along the phylogeny under an HKY substitution model with the transition-transversion ratio $\kappa$ drawn from an exponential distribution with rate 0.15, consistent with estimated values for mitochondrial control regions in Barker et al. (2012).

*S1.5 Details of molecular dating analyses*

In BEAST, we set the rate prior to the uncorrelated lognormal clock with all default parameters. For PAML (mcmctree), the clock is set to the autocorrelated lognormal clock (clock = 3) or uncorrelated lognormal clock (clock=2) with the conditional i.i.d prior with a mean of 0.5 per 10 million years and a single gene. Rate priors and calibrations were scaled to operate in time units of 10 million years, as suggested by the PAML documentation. In addition to the rate prior, we must also specify the prior on tree topologies and/or node times, which represents our prior beliefs about the distribution of node times in the absence of sequence data or calibrations. We assume that the hypothetical researchers carrying out the analysis do not have any prior knowledge of the speciation parameters. For all analyses, we therefore assign a birth-death prior on node times with a broad Uniform (0, 1000) prior on speciation rates, which is the BEAST default with a finite upper bound. All other parameters were left as default settings. For the birth-death prior in PAML (mcmctree), we cannot set hyperpriors, and so we set the birth-death parameters to the true ancestral values at 0.12 and 0.0045. For both BEAST and PAML, Markov Chain Monte Carlo (MCMC) was conducted until an effective sample size (ESS) of at least 100 was reached for all parameters. In molecular dating a threshold of 200 is the normal standard, but we apply a lower standard because our simulations deliberately violate the assumptions of the inference methods leading to very long convergence and mixing times. We assessed mixing visually using the software

Tracer (Rambaut et al. 2018). Highest Posterior Density intervals (HPDs) and median node times for BEAST were calculated using the program TreeAnnotator.

**Table S1.** Analyses used to find a plausible value for the covariance between substitution rates and speciation rates. We simulated data sets from a theoretical correlation of 0.2 and doubled the covariance until a majority of simulated data sets (n = 10, 50 trees each) showed a significant correlation in a mock sister pairs test. For the sister pairs tests, simulated trees were divided at the root to form sister pairs. Contrasts between the clade sizes of the two halves were then regressed through the origin against contrasts between average branch lengths of the two halves. Also shown are the number of individual trees with significant correlations between the sum of branch lengths and number of speciation nodes on root-to-tip paths using the test of Webster et al. (2003) and Pagel et al. (2006). The latter study found 35 ± 4.8% of trees with a significant relationship.

| **Instantaneous covariance** | **Theoretical correlation** | **Sets of 50 trees with significant sister pair correlations (total sets = 10)** | **Individual trees with significant pathlength vs. node number (total = 500)** |
|---|---|---|---|
| 0.0011 | 0.2 | 1 | 43 (8.6%) |
| 0.0022 | 0.4 | 3 | 60 (12%) |
| 0.0044 | 0.8 | 9 | 176 (35.2%) |

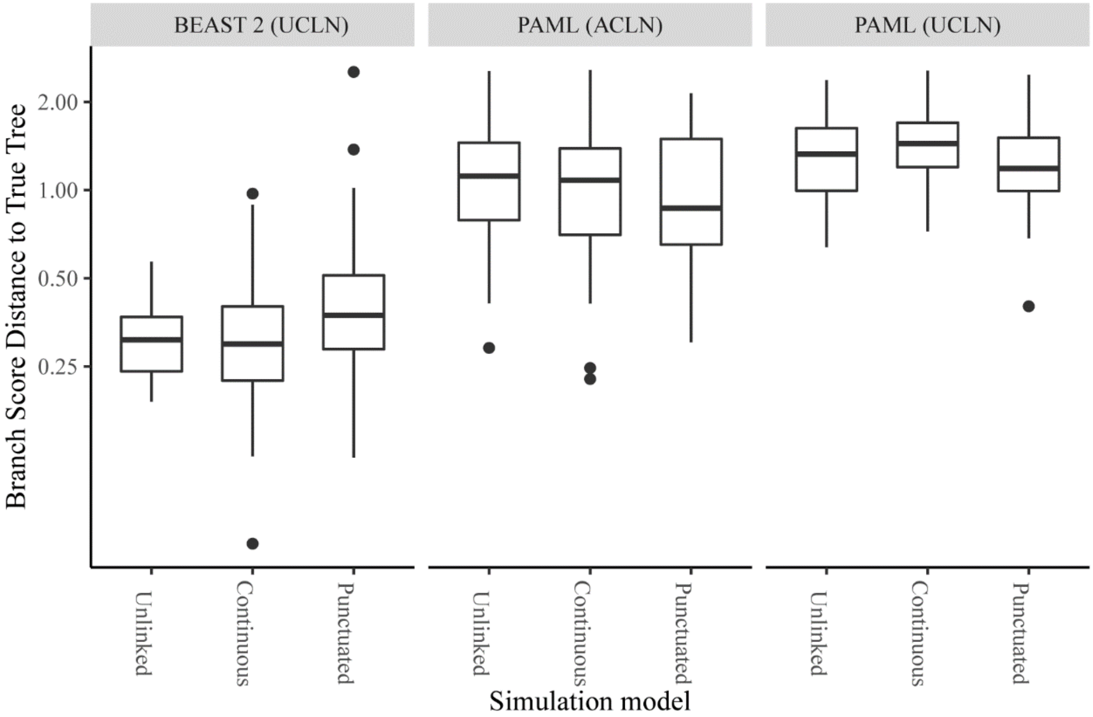


**Figure S1.** Branch score distances from reconstructed to true trees. The median absolute percentage error of node times (MAPE) metric used in the main text is calculated only for branches whose bipartitions are shared between the true and inferred trees, and therefore ignores errors in topology which may occur in the BEAST 2 analyses. Here we calculate an alternative metric using the branch score (Kühner-Felsenstein) distance to the true tree. This distance includes all nodes and sets branch lengths not shared between the two trees to zero, thus reasonably incorporating topological errors (Kuhner and Felsenstein 1994). The three simulation models are Unlinked (instantaneous covariance of molecular rates and speciation rates = 0), Continuous (instantaneous covariance = 0.0044), and Punctuated (instantaneous covariance = 0, bursts of substitutions added at speciation events). Topologies and node times were reconstructed using three different analytical methods, with an uncorrelated lognormal 'relaxed clock' rate prior (UCLN) in BEAST 2, the autocorrelated lognormal rate prior (ACLN) in PAML, and the UCLN in PAML.
